# EF-P-resolved ribosome stalling and collision trigger Rae1-mediated mRNA decay

**DOI:** 10.64898/2026.10.09.757949

**Authors:** Alexandre D’Halluin, Hannah J. LeBlanc, Laëtitia Gilet, Ramy Kazan, Carine Tisné, Gene-Wei Li, Ciarán Condon, Frédérique Braun

**Author notes:** These authors contributed equally and are listed alphabetically (Alexandre D’Halluin, Hannah J. LeBlanc).

## Abstract

Ribosome stalling compromises proteome integrity and cellular fitness, and requires efficient mechanisms to eliminate problematic mRNAs and rescue blocked ribosomes. We previously identified Rae1 as an endoribonuclease that cleaves mRNAs at the tail end of stalled ribosomes to promote mRNA degradation in *Bacillus subtilis*. Here, we show that Rae1 has a broader role in the quality control of mRNAs that encounter translation problems. Global 5′-end sequencing identified 31 previously unknown Rae1 cleavage sites, many of them associated with ribosome pausing at polyproline motifs in the absence of elongation factor P (EF-P). We also demonstrate that Rae1 cleaves mRNAs with ribosomes stalled on truncated transcripts lacking a stop-codon (non-stop mRNAs), provided that the transcript is sufficiently long to accommodate multiple collided ribosomes. Rae1 cleavage depends on a cytosine-poor context at the cut site, showing that both ribosome collisions and RNA sequence at the cleavage site govern Rae1’s ability to promote the removal of stalled ribosomes from mRNA.

**Significance:** Cells have evolved mRNA quality-control mechanisms to identify and eliminate defective messenger RNAs. Ribosome stalling, which occurs when ribosomes encounter difficult-to-translate sequences, can contribute to the production of truncated proteins and compromise cellular fitness. We have identified Rae1 as an endoribonuclease that detects and cleaves mRNAs associated with collided ribosomes, allowing stalled ribosomes to be rescued and transcripts to be eliminated. We show that this mechanism operates broadly across different types of problematic mRNAs and depends on both ribosome collisions and a cytosine poor sequence at the cleavage site. These findings reveal how bacteria couple translation surveillance to mRNA degradation and provide new insights into how cells recruit RNases to promote ribosome rescue and recycling.

## Introduction

Protein synthesis by ribosomes is a costly and time-consuming processes in bacterial cells. Translation efficiency depends, in part, on the ability of the 30S small ribosomal subunit to bind mRNA to the Shine-Dalgarno sequence. However, translation is not always smooth sailing, and ribosome pausing or stalling can impact its efficiency during the elongation step (Buskirk et Green 2017; Samatova et al. 2020; Keiler et Feaga 2014; Buskirk 2026; Takada 2026). Ribosome stalling can be caused by truncated or modified RNA, RNA structure, rare codons, certain ribosome-targeting antibiotics, specific combinations of amino acids such as polyproline motifs or relatively ill-defined ‘arrest peptides’ (Choi et al. 2020; Seip et Innis 2016; Arenz et al. 2014; Chiba et al. 2023; Woolstenhulme et al. 2013; Fostier et al. 2023; Sohmen et al. 2015; Müller et al. 2021; Wilson et al. 2016). The translation elongation factor P (EF-P) has been shown to promote the resumption of translation on polyproline and other stalling motifs (Woolstenhulme et al. 2015; Hersch et al. 2013; Peil et al. 2013; Hummels et Kearns 2020). Recently, the ABCF ATPase YfmR/Uup and YebC2 were shown to play an important supporting role in *Bacillus subtilis*, and loss of these factors incurs a severe fitness defect in the absence of EF-P (Feaga 2026). If left unresolved, ribosome stalling can affect cell viability by dramatically reducing the pool of available ribosomes, or by leading to the production of truncated and potentially toxic proteins. Consequently, bacteria have developed various ribosome rescue mechanisms, notably the trans-translation pathway, driven by the transfer-messenger RNA (tmRNA) in conjunction with small protein B (SmpB), and the back-up rescue systems ArfA and BrfA in *Escherichia coli* and *B. subtilis*, respectively (Keiler 2015; Shimokawa-Chiba et al. 2019; Feaga et al. 2014; Moore et Sauer 2007). *B. subtilis*, also has a distinct rescue pathway that shares similarities with the eukaryotic pathway known as ribosome-associated quality control (RQC) involving ribosome sub-unit splitting and the protein RqcH (Takada et al. 2021; Takada 2026).

Recently, it was demonstrated in *E. coli* that the small MutS-related (SMR) protein, SmrB, is activated by ribosome collisions, which occur when trailing ribosomes catch up to a stalled ribosome, triggering the formation of a specific collided structures in which two adjacent ribosomes lie perpendicular to each other (Cerullo et al. 2022). SmrB cleaves the mRNAs between the stalled and collided ribosome and triggers rescue by the tmRNA pathway (Saito et al. 2022). In *B. subtilis,* the MutS2 protein, which contains an SMR domain, also binds ribosomes at the collision interface but, unlike SmrB, MutS2 does not have RNase activity (Park et al. 2023). Rather, it uses its ATPase activity to split the large and small ribosomal subunits of the stalled ribosome, causing their release from the RNA.

We have recently identified a novel mechanism of mRNA quality control that involves ribosome stalling and mRNA cleavage by the endoribonuclease Rae1 in *B. subtilis* (Lalaouna et Massé 2017; Leroy et al. 2017). Rae1 has no homology to either SmrB or MutS2. We showed that Rae1 cleaves mRNA in a translation-dependent manner and promotes ribosome rescue by the tmRNA, similar to SmrB in *E. coli* (Deves et al. 2023). We identified three mRNAs that are targeted by Rae1 within their open reading frames (ORFs), *spyA* and *bmrX,* encoding a small toxin (17 aa) and a regulatory peptide (26 aa), respectively, and the *fliY* mRNA, encoding the flagellar C-ring protein, the latter in the absence of EF-P (Deves et al. 2025; Hummels et Kearns 2019). Rae1 cleaves these three mRNAs 12 nucleotides (nts) upstream of the A-site of a stalled ribosome with either a stop codon (*spyA, bmrX*) or a proline codon in the A-site (*fliY*). Rae1 cleavage of the *spyA* and *bmrX* mRNAs depends on ribosome stalling, provoked by the interaction between the nascent peptide chain and the exit tunnel of the ribosome, and on the action of release factors (RF2 and RF1, respectively) at their cognate stop codons (Deves et al. 2025). For the *fliY* mRNA, Rae1 cleavage depends on translation of an SPP motif, which causes ribosomes to stall in the absence of EF-P.

Here, we show that Rae1 plays a much broader role in quality control elimination of mRNAs that encounter translation problems than previously understood. A global 5’-end sequencing approach performed in the absence of EF-P allowed us to map 31 novel Rae1 cleavages. We show that Rae1 can also cleave truncated mRNAs lacking a stop codon (non-stop mRNAs) that promote ribosome collisions, which allowed us to analyse cleavage specificity independently of arrest peptide sequence. A mutation analysis of nucleotides around the cleavage site showed that Rae1 has relatively low specificity, but that the presence of cytosine residues in the vicinity of the cut site is correlated with reduced cleavage efficiency. Lastly, we show that open reading frame (ORF) length is important, with at least 15 codons required for efficient cleavage, sufficient to accommodate three collided ribosomes.

## Results

### Identification of 31 novel Rae1 targets by 5’end-seq

We have previously shown that Rae1 promotes the degradation of the *fliY* mRNA in the absence of EF-P (Deves et al. 2025). Cleavage occurred 12 nts upstream of the second proline codon of a known SPP ribosome stalling motif in the *fliY* ORF. To identify novel Rae1 cleavage sites in the *B. subtilis* transcriptome, we performed global 5’-end sequencing at single-nt resolution in the presence and absence of EF-P and/or Rae1. The strains used also lacked the 5’-3’ exonuclease RNase J1 (encoded by the *rnjA* gene) to stabilize the downstream mRNA fragments produced by Rae1. A similar stabilized end-sequencing approach was previously used to globally map cleavage sites of endoribonucleases RNase Y, RNase III and EndoA in *B. subtilis* with single-nt resolution (Taggart et al. 2025).

To assign 5’ ends to Rae1 cleavage events, we compared these putative endonuclease sites, identified computationally using a peak-calling approach described previously (Taggart et al. 2025), in 4 mutant strains: *ΔrnjA, ΔrnjA rae1, ΔrnjA efp* and *ΔrnjA rae1 efp*. The two well-characterized Rae1 sites within the *bmrX* and *spyA* ORFs were captured by our sequencing-based approach, *i.e.* a peak was observed 12 nts from the stop codon in both cases in the *ΔrnjA* strain, but not in the *ΔrnjA rae1* double mutant (Fig. 1A) (Deves et al. 2023; Leroy et al. 2017). An additional Rae1 site was identified with this pair of strains within the *pupG* ORF of the *drm-pupG* bicistronic mRNA, encoding purine nucleoside phosphorylase (Fig 1B) (Table 1). The PupG amino acid sequence did not contain any features known to promote ribosome pausing and the nucleotide sequence surrounding the Rae1 cleavage site showed no obvious similarities with the other known Rae1 sites. The stability of the *pupG* mRNA was slightly increased in the *Δrae1* strain (1.3x fold-change (FC)), suggesting that cleavage by Rae1 accelerates its degradation, albeit to a small degree (Fig. 1C).

**Figure 1:**
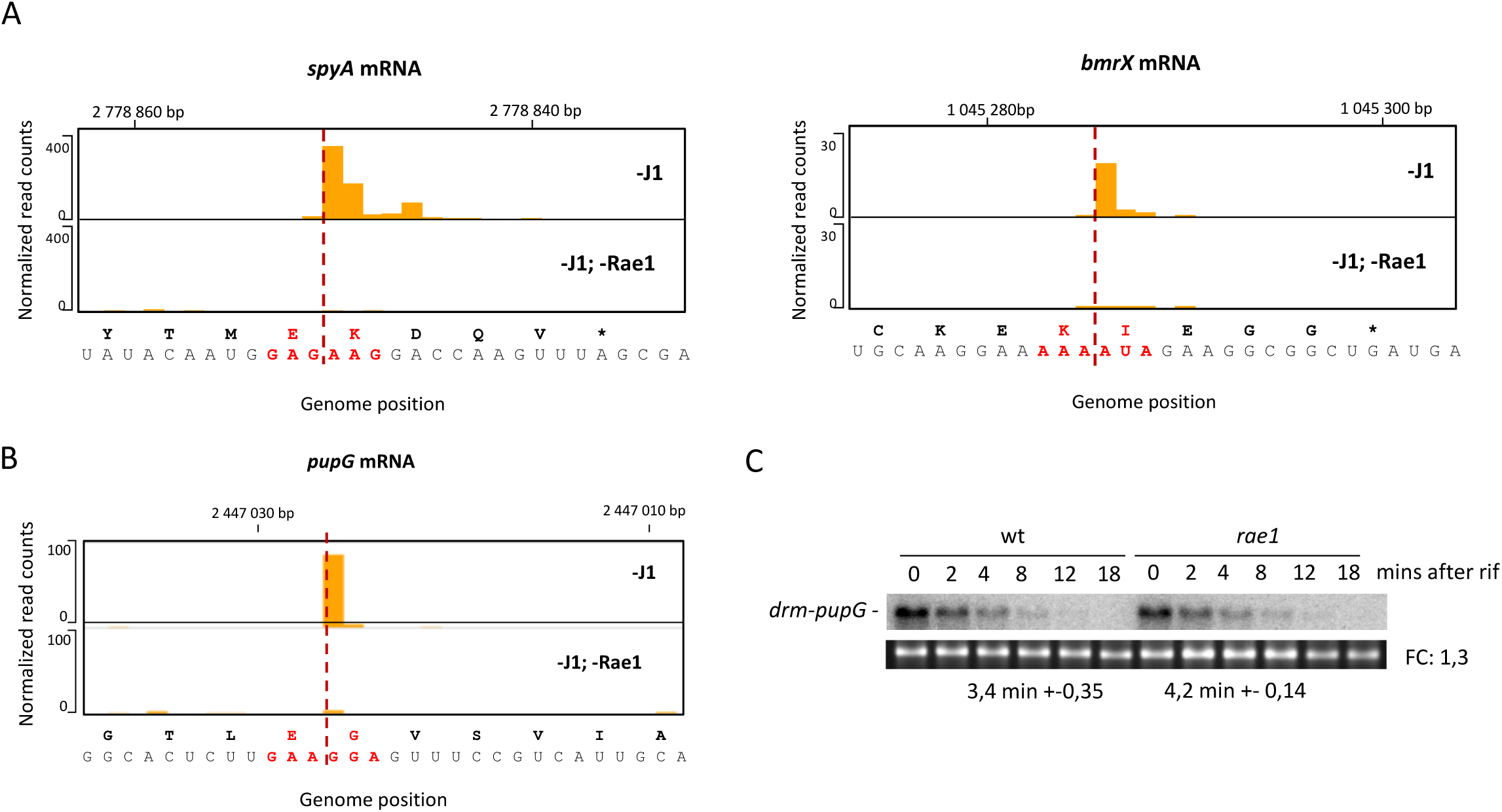
**Rae1 cleavages within *spyA*, *bmrX* and *pupG* mRNAs identified by 5’end-seq.** 5’end-seq profile of (A) the known substrates *spyA* and *bmrX* mRNA, (B) and the newly identified *pupG* mRNA comparing a △*rnjA* strain and a △*rnjA rae1* strain. The Rae1 cleavage site mapped between the two codons (red) is indicated by a red line. The black bar represents the normalized read counts scale. (C) Stability of *pupG* mRNA revealed by northern blotting of total RNA at times after rifampicin addition. The half-life of *pupG* mRNA in the wild-type (wt) and the △*rae1* deletion mutant was calculated using 2 biological replicates. The RNAs were probed using oligonucleotide CC3514. 16S RNA is shown as a loading control and was revealed using 1/20000 Gel Red staining on the agarose before transfer of the total RNA to the membrane.

**Table 1:**
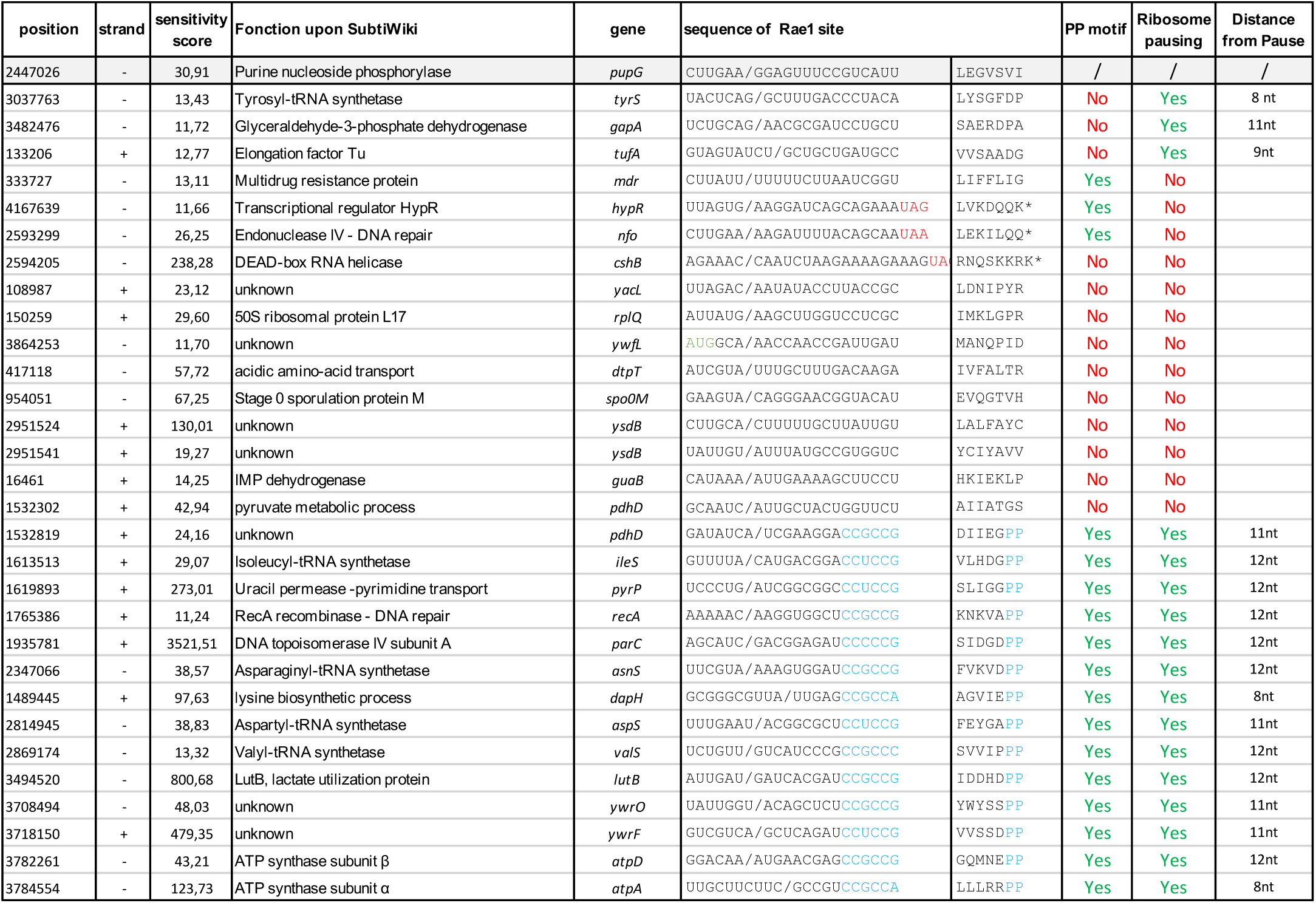
Identification of novel Rae1 targets by 5’end-seq. The sequence and the location of Rae1 cleavage sites are indicated. Prediction of ribosome pausing using Hummels *et al*., 2019 (Hummels et Kearns 2019) ribosome footprinting processed data is indicated by YES or NO, as well as the distance between the pause and Rae1 cleavage site. A score of at least 2 in the △*efp*/wt coverage of was considered as a pause.

A comparison of peaks in the *ΔrnjA efp* and *Δ rnjA efp rae1* strains led to the identification of a further 30 novel Rae1 cleavage sites (Fig. 2A and Table 1). None of these Rae1 cleavage sites were detected in the *ΔrnjA efp*^+^ strains, indicating that they were all EF-P dependent (Fig 2A). The *ysdB* and *phdD* mRNAs, linked to heat stress survival, and to glycolysis and the TCA cycle, respectively, each contained two EF-P-dependent Rae1 cleavage sites. The long polycistronic *atpBEFHAGDC* mRNA involved in ATP synthesis, had an EF-P-dependent Rae1 site in two of its cistrons, *atpA* and *atpD*. Similarly, a Rae1 site was found in both cistrons of the *nfo-cshB* mRNA (Nfo is involved in DNA damage repair in spores, while *cshB* encodes an RNA helicase).

**Figure 2:**
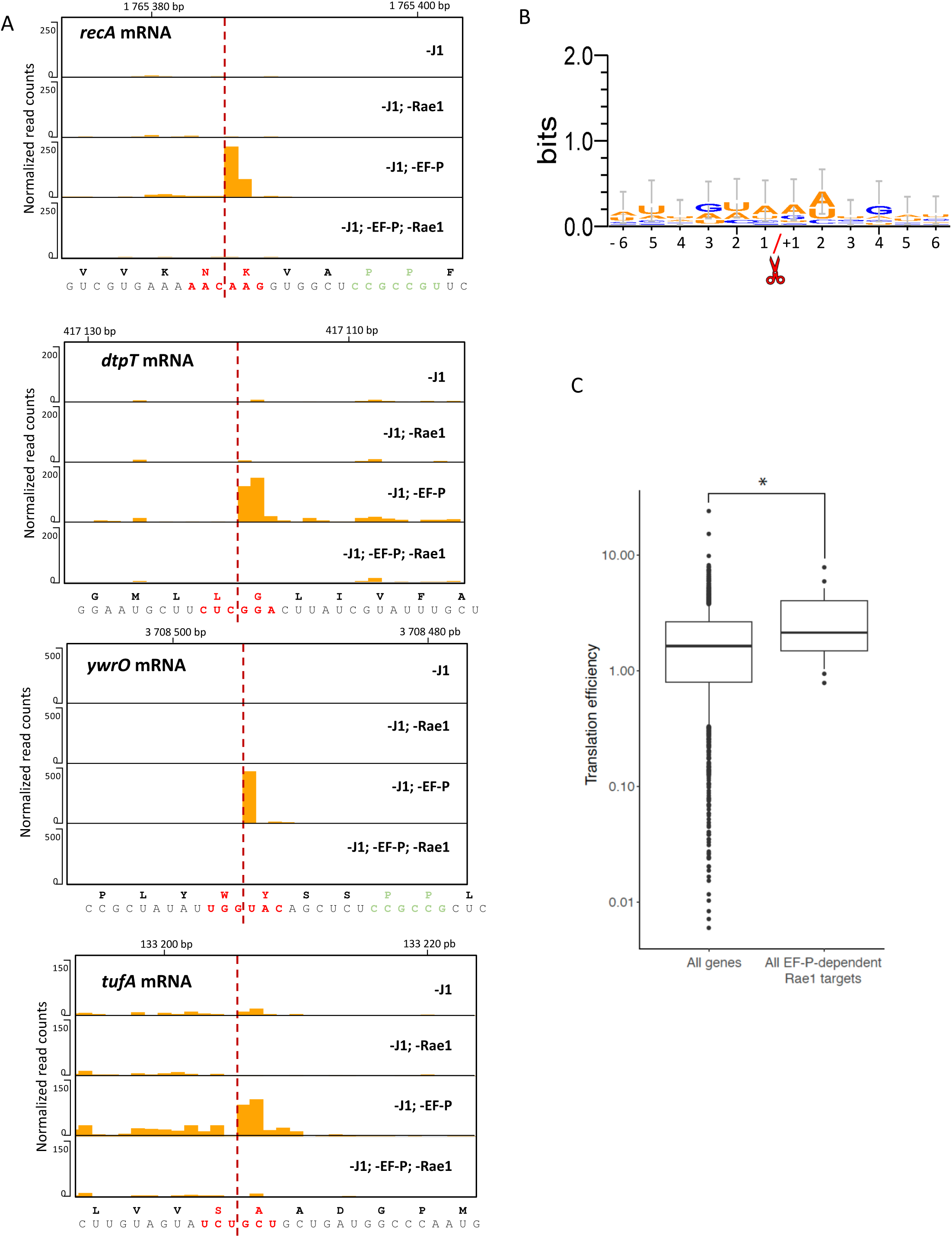
Rae1 cleavage occurs at in a loosely defined sequence context and is related to translation efficiency. (A) 5’end-seq profile of selected regions from *recA*, *dtpT*, *ywrO* and *tufA* mRNA comparing a △*rnjA* strain (J1^-^), a △*rnjA rae1* strain (J1^-^; Rae1^-^), a △*rnjA efp* strain (J1^-^; EF-P^-^), and a △*rnjA efp rae1* strain (J1^-^; EF-P^-^; Rae1^-^). The red line represents the Rae1 cleavage site mapped between the two codons (red). Doublets of proline codons localized 12 nts downstream of the cleavage site are highlighted in green. The left black bar represents the normalized read counts scale. (B) Nucleotide frequency of mapped Rae1 cleavage sites. DNA sequence was extracted from 6 nts upstream and downstream of the mapped cleavage sites from the 31 new targets and the previously characterized *spyA*, *bmrX* and *fliY* sequences. The logo was generated using WebLogo3. Orange: Adenine and Uracyl. Blue: Cystosine and Guanine. (C) Translation efficiency of EF-P dependent Rae1 targets. All genes included in analysis have an abundance of at least 40 transcripts per million as calculated by Rend-seq 3’-mapping reads in at least one of the two Δ*efp* strains, chosen as the observed lower limit of expression for detecting a Rae1 signal. Boxes indicate 25th to 75th percentile; whiskers indicate 10th and 90th percentiles, individual points indicate genes in the 5th and 95th percentiles. * Wilcoxon rank sum p < 0.05 (p = 0.0291)

A polyproline motif was located just downstream of the 5’-end peak in 14 of the 30 mRNAs that contained an EF-P-dependent Rae1 cleavage site. In all 14 of these cases, a ribosome pause was previously identified at the polyproline motif in a global ribosome profiling experiment performed in a Δ*efp* strain (Hummels et Kearns 2019) (Table 1). Rae1 cleaved these 14 mRNAs between 8 and 12 nts upstream of the second proline codon, predicted to be in the A-site of the paused ribosome (Table 1). The three previously characterized Rae1 cleavage sites all occurred 12 nts upstream of the A-site (Deves et al. 2025). Thus, either trimming of some 5’ ends can occur even in the absence of RNase J1 *in vivo*, or Rae1 has some flexibility in terms of its exact cleavage position relative to the A-site.

The remaining 16 mRNAs with EF-P-dependent Rae1 cleavage site did not have polyproline or a recognizable alternative pausing motif downstream of the 5’-end peaks. For three of these, *tufA*, *gapA* and *tyrS,* a ribosome pause was previously seen 8 to 11 nts downstream (Table 1) (Hummels et Kearns 2019). This is consistent with the idea that EF-P can also prevent ribosome stalling at sequences other than polyproline motifs (Hersch et al. 2013). An EF-P-dependent ribosome pause was not detected for the 13 remaining mRNAs (Hummels et Kearns 2019). It is possible that strain differences (168 vs 3610) or differences in culture conditions explain why EF-P-dependent pauses were not detected at these cleavage sites in the previous study.

### Rae1 cleavage occurs at a loosely defined sequence and is related to translation efficiency

Deletion of *efp* lead to increased ribosome pausing in 180 mRNAs (Hummels et Kearns 2019), but the majority of these did not contain Rae1-dependent 5’-end peaks. This is consistent with our model that ribosome stalling *per se* is not sufficient for Rae1 cleavage, but that the RNase additionally has a sequence preference around the cleavage site. We generated a sequence logo from the 6 nts on either side of the cleavage site in the 31 novel Rae1 targets and the 3 well-characterized Rae1 substrates (*spyA*, *bmrX* and *fliY)* (Leroy et al. 2017; Deves et al. 2023, 2025). The logo suggests a weak preference for A/U in position +2 relative to the cleavage site and an even weaker bias towards GUAA from position −3 to +1 (Fig. 2B). More remarkable was the fact that the context around the cleavage site lacked C-residues.

Since Rae1 cleavage is translation-dependent, we wondered whether Rae1’s ability to cleave mRNAs in the absence of EF-P was related to translation efficiency. Moreover, it was shown that EF-P dependence only occurs when a given polyproline motif affects the rate of synthesis and that strongly expressed proteins exhibit the greatest dependence on EF-P (Hersch et al. 2014). Using previously published translation efficiency (TE) values for all *B. subtilis* genes (Lalanne et al. 2018), we compared the distribution of TEs of mRNAs targeted by Rae1 in the absence of EF-P compared to the average for all mRNAs. Our results showed that Rae1-targeted mRNAs had higher mean TEs than non-targets (Fig. 2C). This supports our model that ribosome collisions, which would be predicted to occur more frequently at ribosomes stalled on more heavily translated mRNAs, are the targets for Rae1 recognition (Deves et al. 2025).

### Rae1 targets are generally stabilized in the *Δefp rae1* strain

To validate the novel Rae1 targets identified by stabilized end-sequencing, we asked whether their degradation was Rae1-dependent *in vivo*, by measuring their mRNA half-lives in *Δefp* and *Δefp Δrae1* strains by Northern blot. Among the eight mRNAs tested, seven (*recA, ywrO, pdhABCD, tufA, ileS, asnS, valS*) were stabilized between 1.3 and 2.5-fold in the *Δefp Δrae1* double mutant strain compared to the *Δefp* mutant strain, suggesting that Rae1 indeed catalyzes a rate-limiting step in the degradation of these mRNAs (Fig. 3). The stability of the *dtpT* mRNA, which encodes a peptide transporter, remained unchanged upon *rae1* deletion, suggesting that, in this case, Rae1 cleavage is not rate-limiting for degradation (Fig. 3).

**Figure 3:**
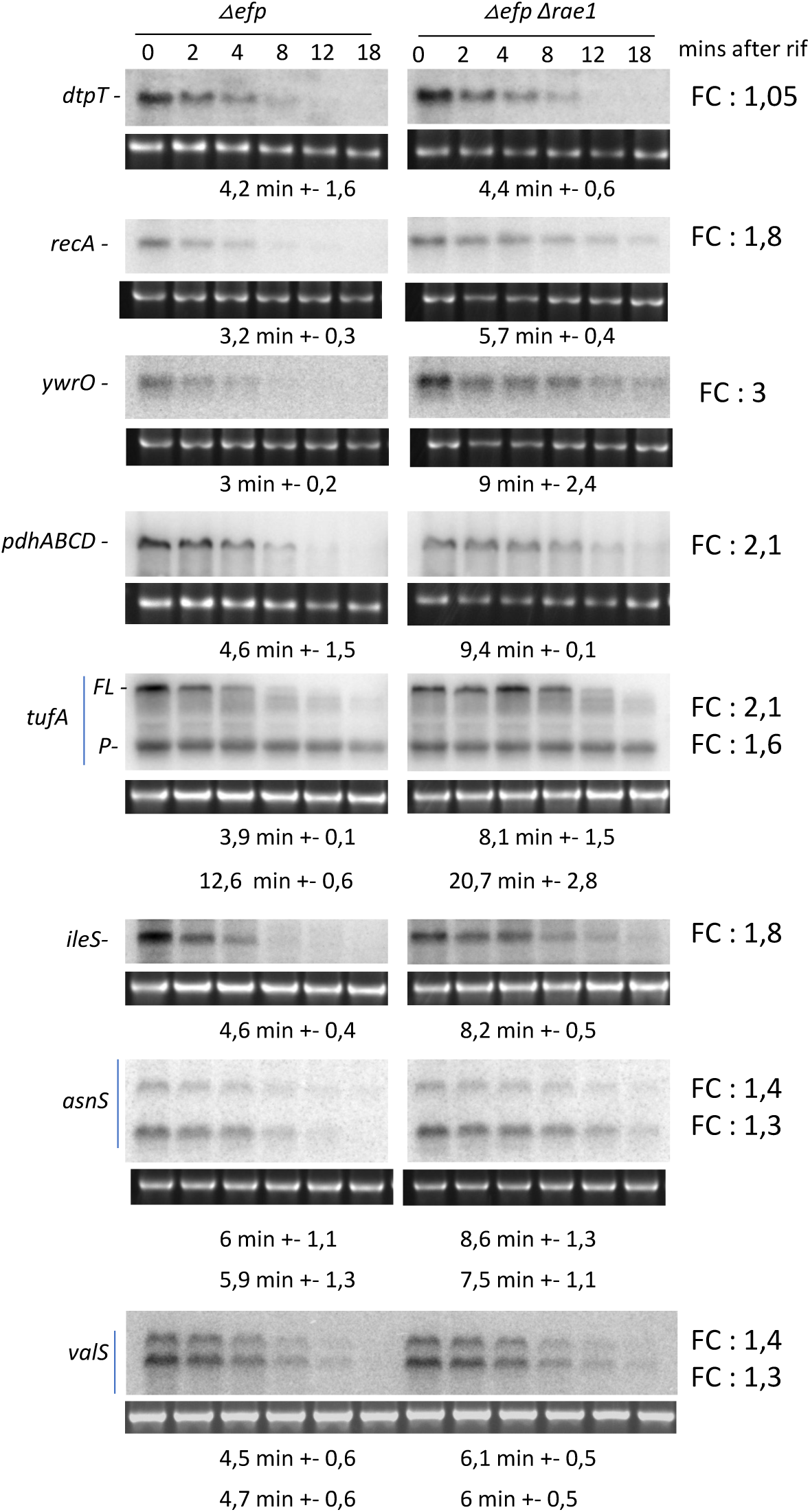
The novel EF-P dependent Rae1 targets are stabilized in a △*rae1 efp* strain. The half-lives of 8 mRNAs identified in the 5’end-seq experiment were determined using 2 biological replicates. 16S RNA is shown as a loading control and was revealed using 1/20000 Gel Red staining on the agarose before transfer of the total RNA to the membrane. RNAs were probed using primers described in Supplementary Table 1. The representative northern blots of the *dtpT* and *pdhABCD* mRNAs, of *recA* and *ywrO* mRNAs, and of *tufA* and *ileS* mRNAs were probed respectively on the same northern blots. The fold-change (FC) in half-life is given next to each blot. FL= Full Length mRNA. P= Processed mRNA.

YfmR is a member of the ABC-F ATPases family that, like EF-P, interacts with the E-site and resolves ribosome stalling at polyprolines (Hong et al. 2024). We thus investigated whether deletion of *yfmR* also had an impact on the stability of the novel Rae1 targets. However, the stability of all 8 mRNAs is similar in *ΔyfmR vs ΔyfmR rae1* strains (Fig. Sup 1), indicating that that these targets are primarily dependent on EF-P to resolve stalling.

### A polyproline pausing motif restores Rae1 cleavage of the *spyA-AAmod* transcript

We previously showed that a modified version of the *spyA* mRNA, in which the reading frame was shifted for all but three codons (EKD) around the cleavage site (*spyA-AAMod),* was no longer cleaved by Rae1 *in vivo*, despite the fact that the native cleavage site was maintained 12 nts upstream of the stop codon (Deves et al. 2025). These experiments showed that the native *spyA* ORF codes for a ribosome arrest peptide and that in the absence of pausing, the cleavage site alone is not sufficient to promote cleavage. To provide further support for the link between polyproline stall sequences and Rae1 cleavage, we asked whether it was possible to restore Rae1 cleavage of the *spyA-AAMod* construct, by introducing an SPP or PP motif at the appropriate distance downstream of the cleavage site. These constructs were called *spyA-AAModSPP* and *spyA-AAModPP*, respectively (Fig 4A). The ability of Rae1 to cleave these two transcripts was evaluated in coupled *in vitro* translation/cleavage (IVT/C) reactions primed with *B. subtilis* ribosomes in the presence or absence of Rae1. As previously observed, a Rae1 cleavage product of the expected size (72 nts) was observed with the WT *spyA* construct but not with *spyA-AAMod* (Fig. 4B). In contrast, the *spyA-AAModSPP* and the *spyA-AAModPP* transcripts were both cleaved by Rae1 *in vitro*, albeit about half as efficiently as WT (13% and 16% efficiency, respectively, vs 28%). The cleavage product from the *spyA-AAModPP* construct was about 3 nts shorter than that from the *spyA-AAModSPP* transcript, suggesting that Rae1 cleavage was shifted from the native cleavage site to maintain the 12-nt distance to the second proline codon in the A-site (Fig. 4A and 4B).

**Figure 4:**
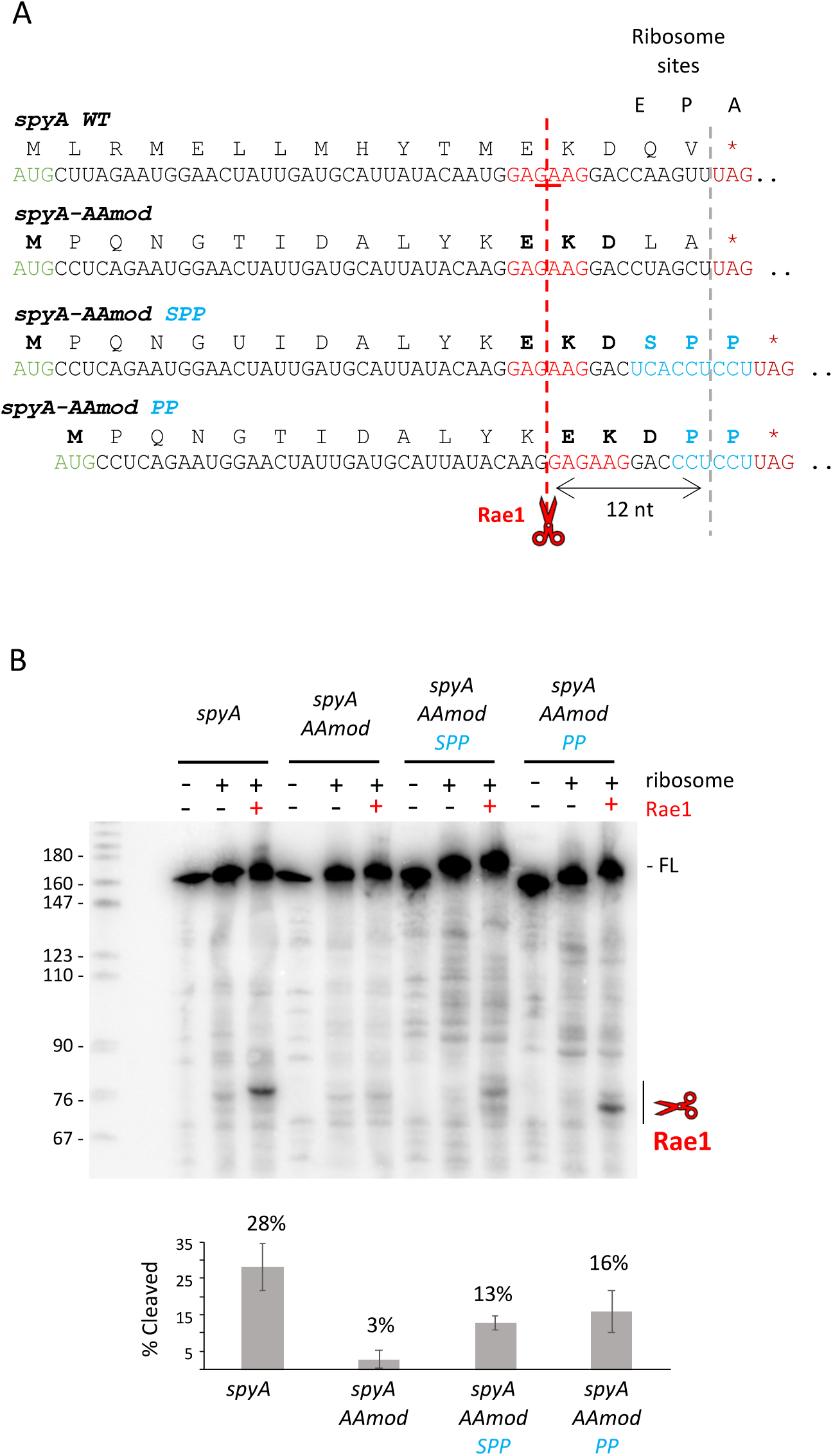
A polyproline stalling motif restores Rae1 cleavage of the *spyA-AAmod* transcript *in vitro*. (A). Sequence of the *spyA WT*, −*AAmod*, −*AAmod SPP* and -*AAmod PP* mRNAs. The Rae1 cleavage site (red bar) is located 12 nts upstream of the A-site of a stalled ribosome. The known and predicted cleavage sites on *spyA* mRNAs are indicated in red on the RNA sequence, and in bold on the amino acid sequence. The insertion of the SPP motif restores the cleavage at the known cleavage site of *spyA*, while the insertion of the PP motif changes the cleavage position to 3 nts upstream. The start codon is indicated in green. (B) *In vitro* cleavage assay of the *spyA* constructs during translation. DNA templates for *in vitro* transcription were generated by PCR using the oligonucleotide pair CC1660/CC1661 on constructed plasmids as described in Material and Methods section. One pmol of *spyA*, *spyA*-*AAmod*, *spyA*-*AAmod SPP* and *spyA*-*AAmod PP* were added to translation assays with 2.7 pmol of ribosomes and 38 pmol of Rae1. A control without ribosomes and Rae1 was included for RNA integrity. The cleaved RNA is indicated by the red scissors. Cleavage assays were done at least in duplicate. FL: Full-length RNA. The proportion of cleaved mRNA was quantified using Fiji software from the signal corresponding to the cleaved band divided by the signal of the cleaved + FL bands.

Together, these data indicate that the introduction of a polyproline pausing motif to an mRNA with a cleavage site 12 nts upstream is sufficient to create a substrate for Rae1 and that the cleavage site can be shifted if an alternative site is available.

### Non-stop mRNAs bearing collided ribosomes are efficiently cleaved by Rae1 *in vitro*

We have shown that the arrest peptides encoded by *spyA* and *bmrX*, and polyproline motifs can promote Rae1 cleavage upstream of stalled ribosomes. We have also found that translation efficiency (this study) and a minimal ORF length that could theoretically accommodate up to three collided ribosomes (Deves et al. 2025), are determinants of Rae1 cleavage efficiency. This led us to propose a model that Rae1 recognizes collided ribosomes. To get further evidence in favor of this model, we decided to force ribosome collisions independently of arrest peptides or polyproline pausing motifs by using a *spyA* mRNA truncated just before stop codon (*spyA-T*) in coupled IVT/C reactions (Fig 5A). First, we showed that the *spyA-T* transcript could provoke ribosome collisions by examining the *in vitro* translation reaction by cryo-electron microscopy (cryoEM). Indeed, the characteristic pattern of perpendicularly-oriented collided disomes and trisomes (Cerullo et al. 2022; Saito et al. 2022) were clearly visible in these translation reactions with the truncated non-stop mRNA (Fig. 5B).

**Figure 5:**
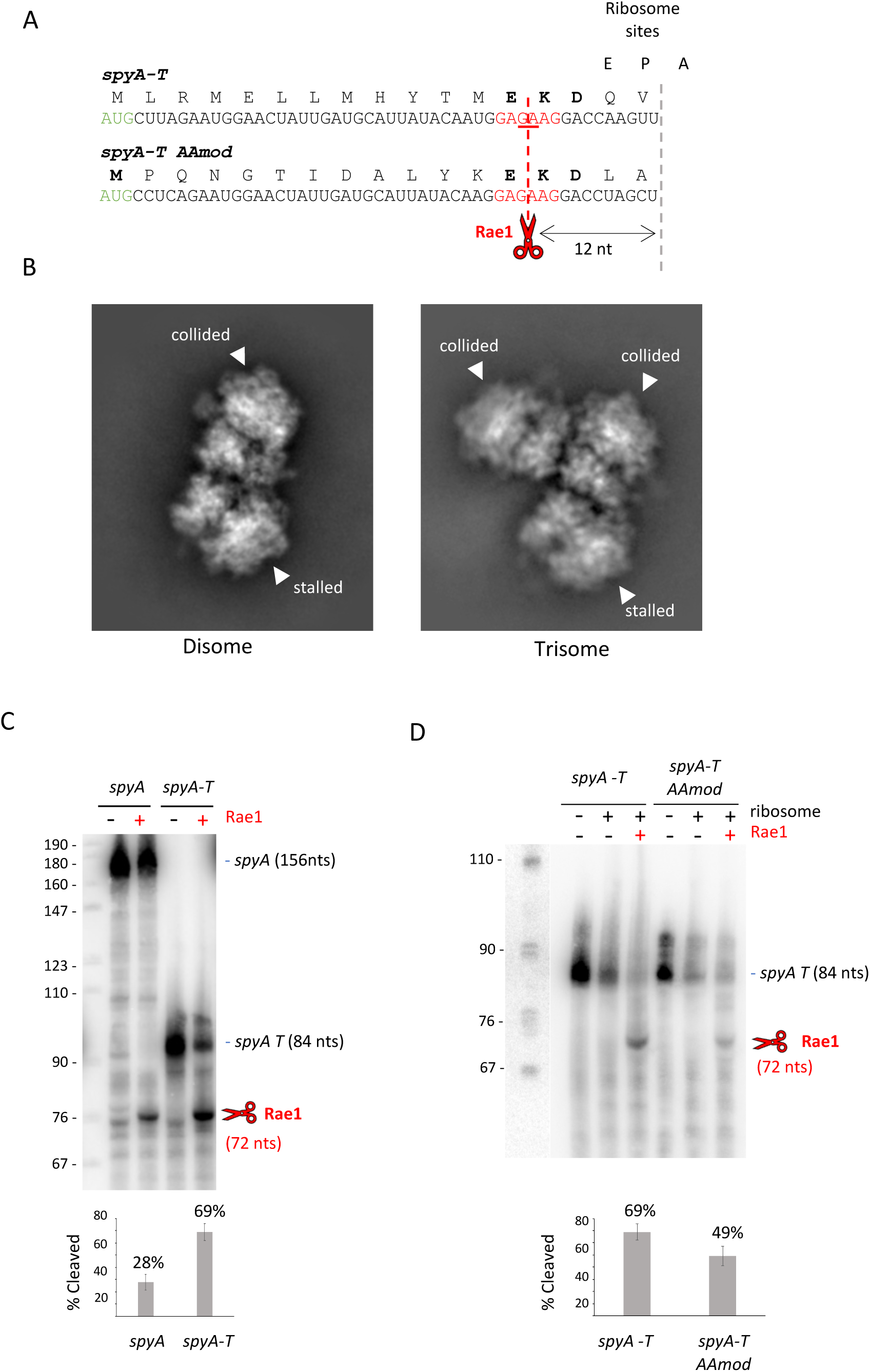
Non-stop mRNAs bearing collided ribosomes are very efficiently cleaved by Rae1 *in vitro*. (A) Sequence of truncated *spyA-T* and *spyA-T*-*AAmod* mRNAs. Rae1 cleavage (red bar) occurs 12 nts upstream of a stalled ribosome with an empty A-site on the truncated mRNA lacking a stop codon of *spyA-T* and *spyA-T*-*AAmod* mRNAs. The start codon is indicated in green. (B) Cryo-EM particles showing disome (left) and trisome (right) formation in the *in vitro* translation reactions of the *spyA-T* transcript. (C and D) Coupled *in vitro* translation and cleavage assay of *spyA, spyA-T* and *spyA-T-AAmod* mRNAs. DNA templates for *in vitro* transcription were generated by PCR using oligonucleotide pair CC1660/CC1661 on constructed plasmids for *spyA* mRNA, or by oligonucleotide hybridization for *spyA-T* and *spyA-T-AAmod* mRNAs as described in the Material and Methods section. The cleaved band is indicated in red. Cleavage assays were done at least in duplicate. The proportion of cleaved mRNA was quantified using Fiji software from the signal corresponding to the cleaved band divided by the signal of the cleaved + FL bands.

We next determined whether Rae1 could cleave the *spyA-T* mRNA at the anticipated site 12 nts from the 3’ end of the transcript. Indeed, Rae1 cleaved *spyA-T* in the same location with more than twice the efficiency (69% vs 28% efficiency) of the full-length *spyA* mRNA. The greater efficiency may reflect a production of more stable ribosome collisions on truncated mRNAs than those induced by a nascent peptide chain and poor translation termination efficiency. These data strongly supporting our model that Rae1 cleaves transcripts bearing collided ribosomes (Fig. 5C). To show that the activity of Rae1 on the *spyA-T* transcript was independent of the peptide sequence, we assayed its activity on a truncated version of the *spyA-AAMod* transcript (*spyA-T-AAMod;* Fig 5A). While the full-length *spyA-AAMod* transcript with a stop codon was not cleaved by Rae1, as seen above (Fig 4B), the truncated *spyA-T-AAMod* transcript lacking the stop codon was cleaved relatively efficiently compared to *spyA-T* (49% vs 69%), showing that cleavage is dependent on the truncation and not the ORF sequence (Fig. 5D).

### ORF length is important

We next took advantage of the fact that Rae1 cleavage of the truncated *spyA* mRNA was independent of ribosome stalling motifs to get a high-resolution picture of the importance of ORF size alone for Rae1 cleavage in coupled IVT/C reactions. Our previous data, based on translational fusions of portions of the *spyA* ORF downstream of *gfp*, suggested that a substrate/non-substrate transition occurred between 14 and 13 codons *in vivo*, but we could not completely eliminate the possibility that the identity of amino acid coded (E5) at the transition point played a role in ribosome pausing (Deves et al. 2025). To address this question, we made variants of the *spyA-T* mRNA (17 codons) with 16 (*spyA*-*T16*), *15* (*spyA*-*T15*), 14 (*spyA*-*T14*), 13 (*spyA*-*T13*), and 12 (*spyA*-*T12*) codons (Fig. 6A), and assayed their ability to be cleaved by Rae1 *in vitro* (Fig. 6B). Rae1 cleaved the wt *spyA-T, spyA-T16* and *spyA*-*T15* mRNAs with over 60% efficiency, cleaved the *spyA*-*T14* construct with intermediate efficiency (26%) and showed no discernable cleavage of the *spyA*-*T13* or *spyA-T12* transcripts (Fig. 6B). Thus, the shorter the ORF length, the less effective Rae1 is at cleaving mRNA, and 15-codon-long ORF (45 nts) sequence appears the shortest length for maximal cleavage efficiency. Since we can see trisomes on the 17-codon *spyA*-*T* mRNA (Fig. 5B), the transition point between 15 and 14 codons in terms of cleavage efficiency may reflect the ability of the mRNA to no longer accommodate three collided ribosomes.

**Figure 6:**
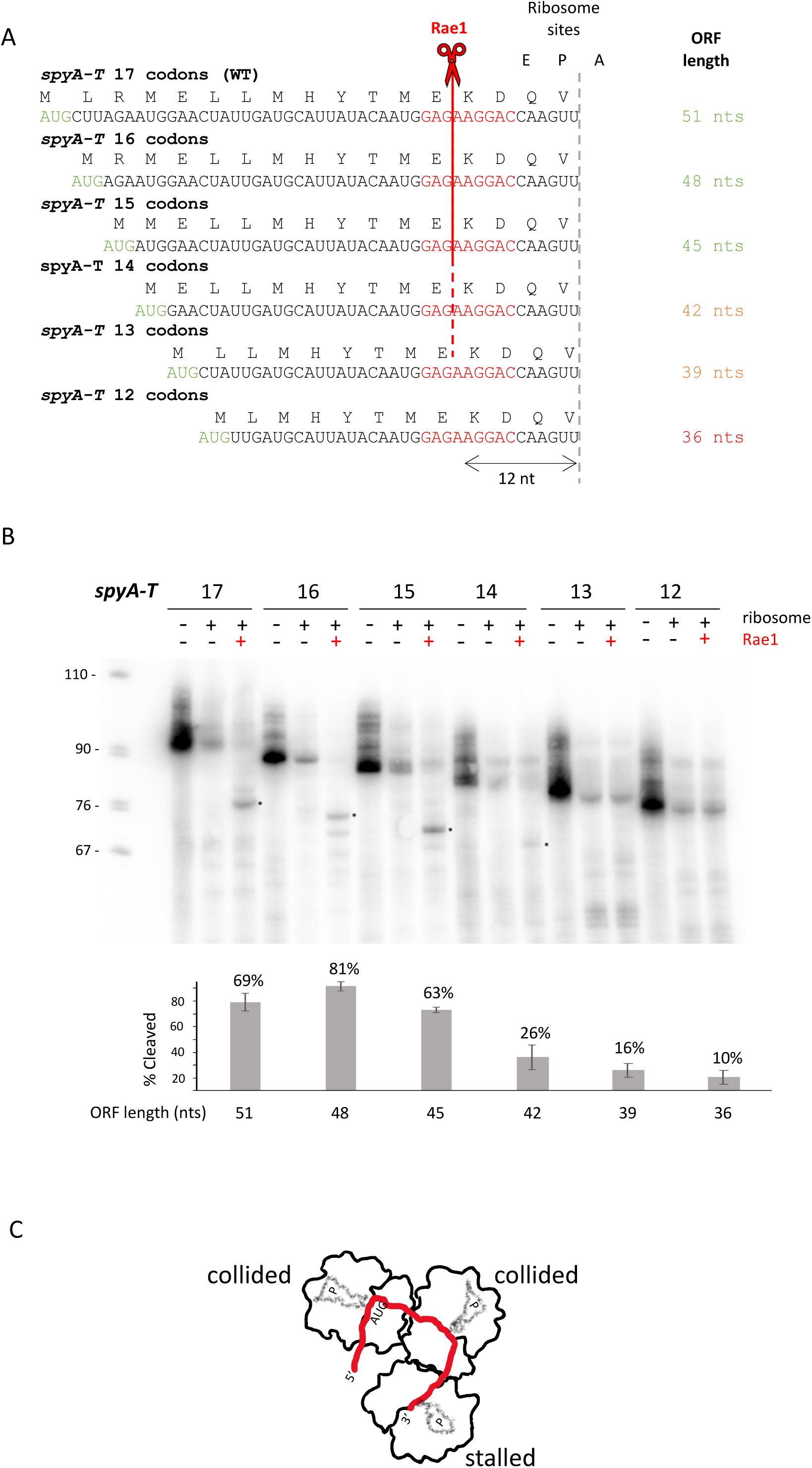
Orf length is important for Rae1 cleavage. (A) Sequence of truncated versions of a *spyA* mRNA with 17, 16, 15, 14, 13 and 12 codons. The position of the full-length mRNA (FL) and the cleaved band are indicated on the right panel. Orf length is indicated. Red bar: Rae1 cleavage site. Green: Start codon. (B) *In vitro* translation and cleavage assay of the truncated versions of *spyA* mRNA. DNA templates for *in vitro* transcription were generated by oligonucleotide hybridization as described in the Material and Methods section. One pmol of mRNA was added to *in vitro* translation reactions with 2.7 pmol of ribosomes and 38 pmol of Rae1. The band corresponding to the cleaved RNA is indicated by a red asterisk. Cleavage assays were done at least in duplicate. The proportion of cleaved mRNA was quantified using Fiji software from the signal corresponding to the cleaved band divided by the signal of the cleaved + FL bands. (C) Schematic of the truncated *spyA-T* mRNA, with two collided ribosomes downstream the stalled ribosome with its empty A-site.

In summary, our data show that Rae1 can cut non-stop mRNAs with ribosomes stalled at their 3′ ends and cleavages typically occur 12 nts upstream of the empty A-site of the stalled ribosome, Moreover, our data suggest that 2-3 collided ribosomes are necessary to induce cleavage, with 3 being better than 2.

### Rae1 does not like cytosine nucleotides

We had previously shown that mutations around the Rae1 cleavage site of the *spyA* mRNA (−6) AUG GAG /AAG GAC (+6) affect the ability of Rae1 to cleave *in vivo* (Leroy et al. 2017). . In particular, the *spyA -3C+1C* mutant spanning the cleavage site and the *spyA +2C* downstream of the cleavage site were stabilized *in vivo* in the *wt* strain, indicating that Rae1 cleavage was strongly reduced (Fig. 7A). The *spyA +5C6U* mutant was also strongly stabilized *in vivo* in the *wt* strain while the *spyA +4A6U* mutant remained unstable similar to WT *spyA* (Fig. Sup2). However, it was impossible to distinguish effects of these mutations on ribosome pausing from effects on Rae1 cleavage preference *in vivo*. Indeed, mutations that do not modify the amino acid sequence, such as the *spyA -1A* and *spyA +3A* mutants, did not interfere with Rae1 cleavage *in vivo* (Fig. 7A) (Leroy et al. 2017).

**Figure 7:**
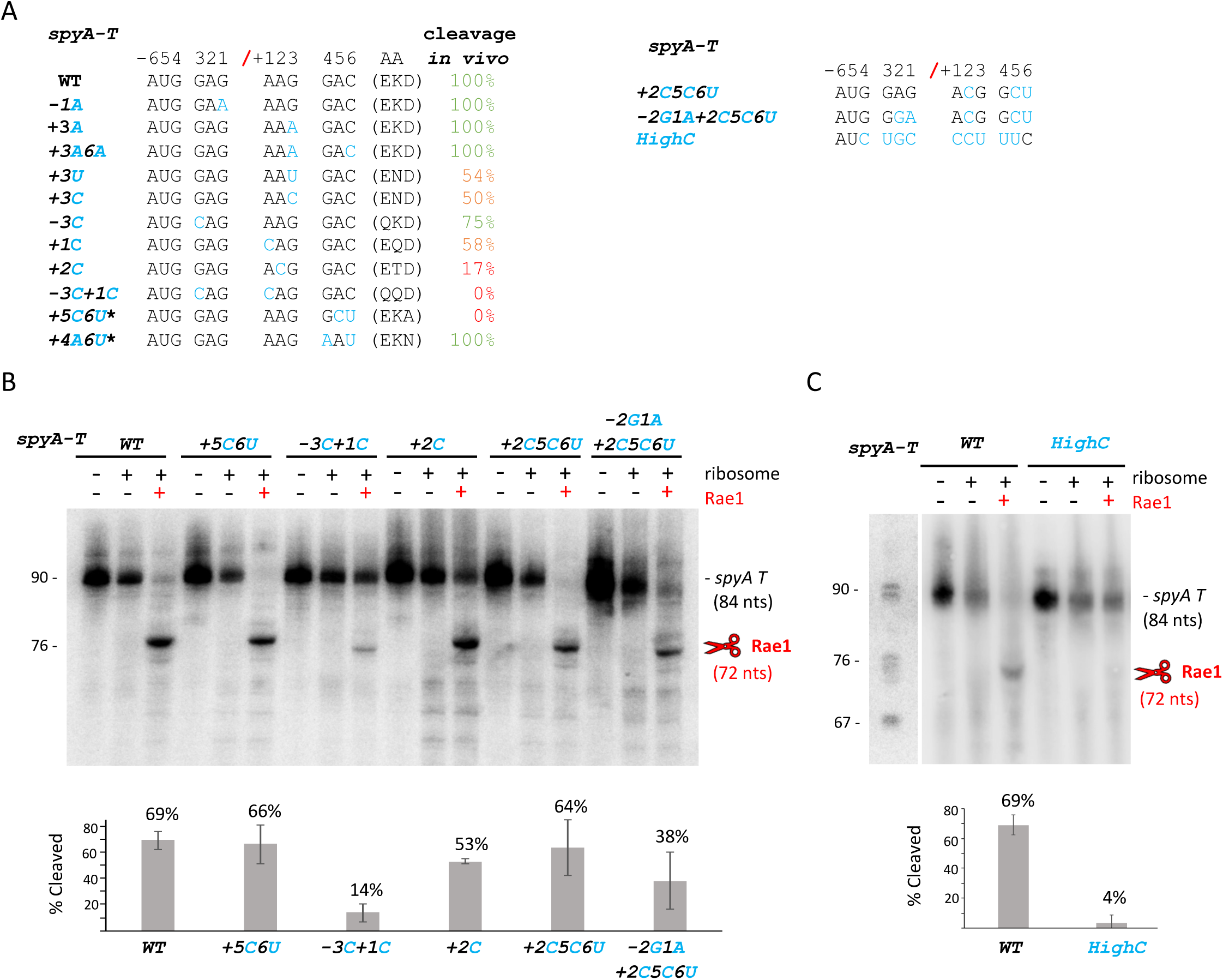
Rae1 does not like cytosine nucleotides. (A) Sequence of the *spyA* mutants. The codons in position 13, 14 and 15 of *spyA* mRNA corresponding to the Rae1 recognition site were mutated at different positions (blue). Positions of the mutations are indicated from the cleavage site position. The amino acid changes from the mutations are indicated next to the nucleotide sequence. The cleavage efficiency *in vivo* from Leroy *et al*., 2017 are indicated on the right panel (Leroy et al. 2017). (B and C) *In vitro* cleavage assay of mutant versions of truncated *spyA-T* mRNA. DNA templates for *in vitro* transcription were generated by PCR or by oligonucleotide hybridization as described in the Material and Methods section. One pmol of mRNA was added to *in vitro* translation reactions with 2.7 pmol of ribosomes and 38 pmol of Rae1. The band corresponding to the cleaved RNA is indicated with red scissors. Cleavage assays were done at least in duplicate. The proportion of cleaved mRNA was quantified using Fiji software from the signal corresponding to the cleaved band divided by the signal of the cleaved + FL bands.

To dissociate the effects of these mutations on arrest peptide sequence from Rae1 cleavage site preference, we tested the ability of Rae1 to cleave mutants showing strong effects *in vivo* in the context of the truncated *spyA-T* construct *in vitro,* which renders RNA cleavage independent of the amino acid sequence (above). The truncated mutant mRNAs *spyA-T -3C+1C, spyA-T +5C6U* and *spyA-T +2C* were synthesized *in vitro* and added to coupled IVT/C in the presence or absence of Rae1 (Fig. 7B). A cleavage product was observed for all three truncated transcripts (Fig 7B). However, cleavage of the *spyA-T -3C+1C* construct was far less efficient (14%) than either *spyA-T +5C6U* (66%) or *spyA-T +2C* (53%). Moreover, the cleaved product of *spyA-T -3C+1C* appeared slightly smaller, suggesting that the cleavage was shifted by a codon to occur between (AUG/CAG CAG). We conclude that the primary effect of the *spyA-T +5C6U* and *spyA-T +2C* mutations *in vivo* is on ribosome pausing due to modification of the amino acid sequence, D15A and K14T, respectively, rather than on Rae1 cleavage site recognition. In contrast, the primary effect of the *-3C+1C* mutation is directly on cleavage efficiency and site selection.

Based on the loose sequence context determined above, which lacked C-nucleotides (Fig 2B), and on the inefficient cleavage of the *spyA-T -3C+1C* transcript, we hypothesized that C-nucleotides might be in general unfavorable for Rae1 cleavage. We there made 3 novel mutants with additional modifications targeting the nucleotides on either side of the cleavage site: *spyA-T +2C5C6U*, *spyA-T -2G1A+2C5C6U* and *spyA-T*-*HighC*. The *spyA-T +2C5C6U* (64%) and *spyA-T -2G1A+2C5C6U* (38%) constructs were both cleaved relatively efficiently (Fig 7B). In contrast, the *spyA-T*-*HighC*, which contained a C at position −4 and 3 consecutive C-residues at the cleavage site (−1 to +2), was barely cleaved (4%) (Fig 7C).

Globally, our data indicate that (i) the A/U preference seen at position +2 in cleavage sites *in vivo* (Fig. 2B) is not critical, as this residue can be mutated to C without a major effect on Rae1 activity *in vitro*, (ii) while the introduction of C nucleotides at positions −3 and +1 individually have only small effects *in vivo*, introduction of both together have a much stronger effect on Rae1 cleavage both *in vivo* and *in vitro,* (iii) in general, the introduction of C nucleotides at or near the cleavage site is correlated with poor cleavage efficiency, albeit not at every position, notably +2 or +5. The fact that certain mutations strongly reduce cleavage efficiency shows that ribosome stalling per se is not sufficient to promote Rae1 cleavage, and Rae1 requires a loose, C-poor sequence motif to cleave 12 nts upstream of the stall site.

## Discussion

Previously, we showed that Rae1 cleaves mRNAs at the tail end of stalled ribosomes, leading to a non-stop ribosome when the next ribosome in the cue moves to the end of the truncated mRNA, allowing it to be rescued by the tmRNA/SmpB (Deves et al. 2023, 2025). Indeed, translation and mRNA decay have been shown to be closely interdependent, ensuring that mRNAs are degraded when they are no longer required for protein synthesis (Deana et Belasco 2005). Prior to this study, only three Rae1 cleavage sites were mapped within the s*pyA* and *bmrX* mRNAs, 12 nts from the stop codon, and within the *fliY* mRNA, 12 nts from the second proline codon of a known stalling motif SPP in the absence of EF-P (Deves et al. 2025). Here, we identified 30 novel Rae1 targets that were cleaved in the absence of EF-P using a 5′ end sequencing approach. Interestingly, several of these EF-P-dependent Rae1 cleavages targeted mRNAs encoding proteins involved in translation: tRNA synthetases (*valS, tyrS, ileS*, *asnS,* and *aspS*), ribosomal protein L17 (*rplQ*) and elongation factor EF-Tu (*tufA*) (Table1). We also demonstrated that Rae1 can cleave truncated mRNAs lacking a stop codon, 12 nts from the 3’ end of the mRNA, thus permitting the clearance of stalled ribosomes from these problematic mRNAs. Together, these findings suggest that Rae1 plays a broad role in ribosome-based mRNA quality control. Supporting these findings, recent work by the Feaga lab has demonstrated that the overexpression of either an ATP hydrolysis deficient ABC-F factor (YdiF(EQ2)) or treatment with erythromycin, both of which cause ribosome collisions, increased tmRNA-mediated tagging *in vivo* that was dependent on Rae1 (Tetreault et al. 2026). In agreement with the authors of this paper, we have named this pathway CART for “**C**ollision-**A**ssociated **R**ae1-induced trans-**T**ranslation”.

Fourteen of the novel EF-P-dependent Rae1 cleavage sites were located upstream of polyproline motifs, consistent with EF-P’s ability to alleviate ribosome pausing on polyproline tracts through its capacity to interact with the CCA of the P-site tRNA^Pro^ (Ude et al. 2013). Ribosome profiling data performed in the absence of EF-P by the Kearns’ lab (Hummels et Kearns 2019) confirmed the presence of a ribosome pause just downstream of the cleavage site for these 14 mRNAs, with the second proline codon of a polyproline motif located in the A-site. For the *tyrS, gapA* and *tufA* mRNAs, Rae1 cleavages also occurred upstream of a ribosome pause site in the absence of EF-P, but not at a polyproline motif: GFD**P**TAD, RDP**A**KLS and AAD**G**PMP, respectively (the amino acid located in the A-site is highlighted in bold). Indeed, almost one third of the ribosome pauses identified in this ribosome profiling experiment (86 out of 246 pauses in 180 different mRNAs detected in the absence of EF-P) were not associated with a polyproline motif (Hummels et Kearns 2019). Consistent with this, it has previously been proposed that EF-P can boost peptide formation between amino acids other than prolines and facilitate the translation of other difficult-to-translate sequences (Hummels et Kearns 2020).

Two additional factors, YfmR and YebC2, have recently been implicated in the resolution of ribosome stalling at polyproline motifs (Hong et al. 2025, 2024; Takada et al. 2024). YfmR belongs to the ABC-F family of ATPases and is closely related to the *E. coli* Uup and EttA proteins (Ousalem et al. 2019). YfmR binds to the E-site of the ribosome and has redundant functions in preventing ribosome stalling at polyprolines and on polyacidic residues (Hong et al. 2024; Takada et al. 2024). YebC2 has been shown to reduce ribosome stalling in the absence of both EF-P and YfmR (Hong et al. 2025). Transcriptome analysis performed in 110 different growth conditions revealed that *yfmR*, *yebC2*, *efp* and *rae1* have remarkably similar expression profiles supporting the idea that they are functionally related (Fig. Sup3) (Nicolas et al. 2012). However, none of the Rae1 targets identified in the absence of EF-P showed altered stability in the absence of YfmR. Further investigations are therefore required to determine the extent to which the absence of YfmR and/or YebC2 can cause ribosome pauses and Rae1 cleavage.

The novel EF-P-dependent Rae1 cleavage sites identified upstream of ribosomes pauses, are located between 8-12 nts from the codon located in the A-site. The cleavage sites of the *spyA, bmrX* and *fliY* mRNAs were all mapped at the tail end of a paused ribosome, 12 nts from a stop codon or a proline codon located in the A site (Deves et al. 2025). In the cases of *spyA* and *bmrX*, ribosome arrest is induced by the sequence of the nascent peptide chain, coupled with the action of release factors at the stop codon, while for the *fliY* mRNA, the ribosome stalls on the second proline codon of a polyproline motif in the absence of EF-P. We showed here that the introduction of a polyproline motif at the appropriate distance from the cleave site in the insensitive *spyA-AAmod* transcript rendered the transcript sensitive to Rae1. Similarly, using the truncated *spyA-T-AAmod* transcript lacking a stop codon to force ribosomes to stall at the 3’ end also restored Rae1 sensitivity to this transcript. The truncated non-stop *spyA* transcript was cleaved less efficiently when only 14 codons remained and was no longer cleaved when the length of the ORF was reduced to 12 codons. Given that there are only about 15 nts (5 codons) from the P-site to the trailing edge of a stalled ribosome (Cerullo et al. 2022), this indicates that a single stalled ribosome is insufficient for cleavage. The fact that a minimal ORF length of 15 codons is required for maximally efficient cleavage by Rae1 suggests that Rae1 is probably preferentially recruited to collided trisomes. Decipher how Rae1 acts more precisely awaits the resolution of the structure of Rae1 in association with collided ribosomes.

The ribonucleases NONU-1 in *Caenorhabditis elegans*, Cue2 in *S. cerevisiae*, and SmrB in *E. coli* use their SMR domains to cleave mRNAs in association with collided ribosomes (Saito et al. 2022; D’Orazio et al. 2019; Glover et al. 2020). Since there is no sequence or structural homology between SmrB proteins and Rae1, the recognition of collided ribosomes by these two different families of RNases appears to be a case of convergent evolution. Interestingly, ORF length is also important for SmrB cleavage. It does not bind to collided ribosomes when the stalling motif is too close to the start codon (Saito et al. 2022), suggesting that multiple ribosomes must also be loaded onto the mRNA for SmrB recruitment. Three collided ribosomes have been suggested to provide a structural basis for preferential Hel2-mediated ubiquitinylation of trisomes compared to disomes, suggesting their architectures are different and this may also provide the basis for discrimination by Rae1 and/or SmrB (Matsuo et al. 2020).

The *B. subtilis* MutS2/RqcU protein, homologous to SmrB, was also shown to bind to collided ribosomes, but it lacks endoribonuclease activity (Park et al. 2023; Cerullo et al. 2022). MutS2 is part of the ribosomal quality control pathway and splits the collided ribosome into 30S and 50S subunits allowing RqcH to promote alanine tagging of the nascent peptide targeting it for degradation (Takada et al. 2021). Thus, *B. subtilis* possess two ribosome rescue pathways, mediated respectively by MutS2/RqcH and Rae1/tmRNA. We previously showed that deletion of *mutS2* had no impact on the degradation of the *spyA* mRNA mediated by Rae1, suggesting that even though MutS2 and Rae1 both recognize collided ribosomes, they act in distinct pathways (Deves et al. 2023). One important determinant of the choice between Muts2 and Rae1 rescue pathways could be the presence of a Rae1 site. Future experiments will be geared towards determining in which conditions and contexts, ribosome collisions recruit MutS2 rather than Rae1, and *vice versa*. In this context, it is interesting to note that a *mutS2* deletion strain is highly sensitive to antibiotics that block translation (erythromycin, chloramphenicol, tetracycline), suggesting that MutS2 intervenes when ribosomes are stalled by antibiotics, while strains lacking Rae1 are more resistant to chloramphenicol (Cerullo et al. 2022; Deves et al. 2023).

Our previous studies have highlighted a role for Rae1 in regulation of expression of the *spyTA* and *bmrBXCD* polycistronic mRNAs, by shutting down the expression of the SpyT toxin (Gilet et al. 2025) and readthrough transcription of the *bmrB* attenuator, which would lead to futile expression of the BmrCD efflux pump in the absence of antibiotics (Deves et al. 2023). In the present study, we demonstrate that the role of Rae1 goes far beyond the regulation of these two transcripts, playing dual roles in gene regulation and translational quality control. Rae1 cleaves multiple mRNAs undergoing translational difficulty, as well as truncated non-stop mRNAs that can occur in cells through physical breakage, premature transcription termination or cleavage by endoribonucleases. Ribosome collisions on these defective mRNA promote Rae1 cleavage to ensure their quality control elimination and the freeing of trapped ribosomes.

## Materials and methods

### Strains and oligonucleotides

Oligonucleotides and strains used in this study are listed in Supplementary Tables 1 and 2 respectively. All *Bacillus subtilis* strains used are derivatives of W168 (Lab strain SSB1002).

The Δharma*efp rnjA* (CCB1538) and the Δ*efp rae1 rnjA* (CCB1555) strains were generated by transformation of the Δ*efp* strain (CCB1481) and Δ*efp rae1* (CCB1481) strains with genomic DNA from a Δ*rnjA* strain (CCB434). The strains were verified by PCR and sequenced.

The spectinomycin resistant (spc^R^) Δ*efp* (CCB1977) strain was constructed by overlapping PCR. The upstream and downstream regions of *efp* were amplified with oligonucleotide pairs CC3596/CC3597 and CC3600/3601 using SSB1002 genomic DNA as template, and the spc^R^ cassette with CC3598/CC3599 oligonucleotides from plasmid pDG1727 (pl1727). The whole construct was amplified using 10 ng of the three PCR products as templates and oligonucleotides CC3596/CC3601. The PCR product was directly transformed into SSB1002 strain. The kanamycin resistant Δ*yfmR* (CCB1979) strain was generated by transforming genomic DNA of strain 168 *trpC2* Δ*efp yfmR* (HAF521) strain, a gift from Heather Feaga, into strain SSB1002. The deletion of *yfmR* was verified by PCR, and the integrity of *efp* locus was verified by PCR and its expression by northern blotting. The Δ*efp rae1* (CCB1978) and Δ*yfmR rae1* (CCB1980) strains were generated by transformation of genomic DNA from Δ*rae1* strain CCB375 into CCB1977 and CCB1979, respectively. All constructions were verified by PCR.

### Plasmid constructions

All plasmids used in this study are listed in Supplementary Table 3. To ensure that the mutated sequences of *spyA* mRNA constructs were present in our *in vitro* Rae1 cleavage assay, DNA templates were cloned in plasmids pBSKS+ or pHM2 and sequence, before amplification by PCR for *in vitro* transcription.

The pHM2-*hbs*Δ-*spyA* (Asp15>Asn) (pl802) and the pHM2-*hbs*Δ-*spyA* (Asp15>Ala) (pl820) plasmids contain the *spyA* ORF with a mutation of codon 15 (GAC to AAU or GAC to GCU), respectively. The mutant *spyA* ORFs were inserted in the 3’ UTR of the internally deleted *Δhbs* gene, described previously (Leroy et al, 2017). The constructs were obtained by overlapping PCR using oligonucleotide pairs CC1607/CC2243 and CC2242/CC572, or CC1607/CC2241 and CC2240/CC572, with pHM2-*hbs*Δ-*spyA* (pl707) as a template (Leroy et al, 2017). The constructs were cloned between the EcoRI and BamHI cloning sites of plasmid pHM2 (pl606). The resulting plasmids pl802 and pl820 were linearized by XbaI and transformed into SSB1002 for integration into the *amyE* locus (Supplementary Table 2), with selection for chloramphenicol resistance, to generate CCB1128 and CCB1153, respectively.

Plasmids pHM2-*spyA-AAmod-SPP* (pl1083) and pHM2-*spyA-AAmod-PP* (pl1087) were made by overlapping PCR using oligonucleotide pairs CC1787/CC3637 and CC3638/CC572 or CC1787/CC3639 and CC3640/CC572, respectively, with pHM2-*spyA-AAmod* (pl1008) as a template (Deves et al, 2025). The overlapping PCR fragment was amplified using 10 ng of PCR products and oligonucleotides CC1787/CC572. The constructs were cloned between EcoRI and BamHI cloning sites of the pHM2 plasmid, as above.

Plasmids pBSKS+*spyA*(GTA) (pl1038) and PBKS+*spyA*(ETA) (pl1039) were generated by amplifying the *spyA* ORF with oligonucleotide pair CC1660/CC3397, to mutate GAGAAGGAC -> GGAACGGCT (EKD->GTA), and with oligonucleotide pair CC1660/CC3396 to mutate GAGAAGGAC -> GAGACGGCT (EKD->GTA). The PCR products were cloned in SmaI linearized the vector pBSKS(+).

### Half-life assay and northern blot

For half-life assays, cells were grown in 2YT medium to OD600 nm = 0.6 and rifampicin was added at 150μg/ml final concentration and harvested 0, 2, 4, 8, 12 and 18 min after addition. Northern blots were performed on total RNA isolated by the RNA-Snap method described in (Stead et al. 2012).

Northern blots were performed as described previously and exposed to PhosphorImager screens (GE Healthcare) (Durand et al. 2012). The signal was obtained by scanning with a Typhoon BioMolecular Imager (Cytiva) and analyzed by Fiji (ImageJ) software.

### Identification of Rae1 cleavage sites by 5’end-seq

Strains were streaked on pre-warmed LB-agar plates and grown for approximately 8 hours at 37°C in 2YT medium. Starter liquid cultures were then inoculated from these plates. To ensure consistent growth across experiments and to minimize lag time, all inoculation after the initial plate streaking was performed at 37°C without removing plates or cultures to lower temperatures. All liquid cultures were vortexed before and after inoculation to prevent clumping. Starter cultures were grown with shaking at 225 rpm for approximately 12 hours, then back-diluted into pre-warmed 2XYT to an OD600nm of 0.0002. Cultures were grown at 37°C with shaking at 225 rpm until OD600 reached 0.2 (i.e. 10 doublings), at which point cells were harvested by combining 7 mL cell culture with 7 mL prechilled 100% methanol. Cells were pelleted by centrifugation at 4000 rpm at 4.C for 10 minutes. Cells were then resuspended in 10 mL pre-chilled 10 mM Tris pH 7 and pelleted again. After removing the supernatant, pellets were flash-frozen in liquid nitrogen and stored at −80°C. For 5’ end-seq, RNA was extracted as described in Taggart et al. 2025 and RNA concentrations were quantified by Qubit RNA BR (Thermo Fisher). gDNA was removed by treating 10 μg total RNA with TURBO DNase (Thermo Fisher) for 1 hour at 37°C and purifying with an RNA Clean and Concentrator-5 kit (Zymo). Purified RNA was eluted in RNase-free water, then brought to 10 mM Tris pH 7 and 1 mM EDTA. For Rend-seq, cell pellets were resuspended by vortexing in 100 μL of lysis buffer (10 mg/mL lysozyme in 10 mM Tris and 1 mM EDTA and incubated for 5 minutes at 37.C. RNA was then purified using the RNeasy Plus Kit (Qiagen) following the manufacturer’s instructions, utilizing the gDNA removal column and adding beta-mercaptoethanol to the RLT-Plus buffer (10 μL BME per 1 mL buffer). RNA was eluted in RNase-free water, then brought to 10 mM Tris pH 7 and 1 mM EDTA.

For both extraction methods, RNA concentration was measured by Qubit RNA BR, DNA depletion was confirmed by Qubit DNA BR, and RNA quality was checked by Fragment Analyzer (Agilent). rRNA was depleted from 10 μg (for 5’ send-seq) or 20 μg (for Rend-seq) of total RNA using the MICROBExpress Bacterial mRNA Enrichment Kit (Thermo Fisher) following the manufacturer’s instructions. 5’ end-seq libraries were prepared and sequenced as described previously, using a post-fragmentation size selection of 60-90 nt (Taggart et al. 2025). Rend-seq libraries were prepared and sequenced as described previously in Taggart et al 2025. Putative Rae1 cleavage sites were identified from 5’ end-seq and Rend-seq by the process described previously using a sensitivity score threshold of 11. The RNA-seq plots were generated using R software. The base-pair probability of Rae1 cleavage site was generated using WebLogo 3 with 6 nucleotides flanking sequences of all 31 newly identified Rae1 cleavage sites and includes the sequence of *spyA*, *bmrX* and *fliY* mRNAs. Previously published translation efficiency values were used for TE comparison (Lalanne et al. 2018), excluding genes with fewer than 40 transcripts per million in 3′-end mapping Rend-seq data from both Δ*efp* strains.

### *In vitro* translation and Rae1 cleavage assay

The *B. subtilis* Rae1 hexahistidine-tagged protein was produced in *E. coli* using the plasmid pET28-Rae1Chis. The 70S ribosomes were purified from 1 g of frozen *B. subtilis* W168 cells as previously described (Leroy et al. 2017).

The full-length *spyA* (pl707), *spyA AAmod* (pl1008), *spyA AAmod SPP* (pl1083) and *spyA AAmod PP* (pl1087) transcripts (Figure 4 and 5) were synthesized *in vitro* from DNA templates generated by PCR with the same oligonucleotides (CC1660/CC1661) and the indicated plasmid.

The truncated *in vitro* transcripts used in Fig. 7 were synthesized from PCR templates amplified using the following oligonucleotides and the indicated plasmid: *spyA* T WT (CC1660/CC3309 and pl707), *spyA* T +2C (CC1660/CC3342 and pl748), *spyA* T +2C5C6U (CC1660/CC3396 and pl1039), *spyA* T -2G1A +2C5C6U (CC1660/CC3397 and pl1038), *spyA* T +5C6U (CC1660/CC3607 and pl741) and *spyA* T -3C+1C (CC1660/CC3606 and pl820).

Truncated *spyA* transcripts were also synthesized from DNA templates obtained by hybridizing two oligonucleotides at 95°C for 5 min, followed by progressive cooling at room temperature in a buffer containing 1 mM EDTA, 50 mM NaCl and 10 mM Trip pH 7.5. For *spyA-T* and *spyA-T AAmod*, the oligonucleotide CC3635 was hybridized with CC3630 or CC3634 respectively. To produce *spyA-T* transcripts shortened in length used in Fig. 6, CC3635 was hybridized with the following complementary oligonucleotides: CC3780 (*spyA*-T (16)), CC3781 (*spyA*-T (15)), CC3631 (*spyA*-T (14)), CC3753 (*spyA*-T (13)), and CC3754 (*spyA*-T (12)). For the *spyA-T HighC* transcript used in Fig. 7C, CC3635 was hybridized with CC3756.

All the transcripts were synthesized by T7 RNA polymerase *in vitro* using a MegaShortScript kit (Ambion) and purified as previously described (Leroy 2017). *In vitro* translation assays (PURExpress <u>Δ</u>ribosomes; New England Biolabs) were typically performed in 5 μl reactions with 1 pmol mRNA, 6.5 pmol *B. subtilis* 70S ribosomes and 38 pmol of Rae1 per reaction. Reactions were incubated at 37°C for 30 min followed by phenol extraction. The aqueous phase was precipitated overnight at −20°C in 3 volumes of ethanol and 1/10 vol of LiCl, supplemented with 1 μl glycogen (20 mg/ml). Pellets were resuspended in 5 μl H2O, with 5 μl RNA loading dye (Biolabs). Six microliters of the samples were run on 8% sequencing gels. RNAs were electro-transferred overnight at 4°C at 15V to a nylon membrane, hybridized using 10ml of ROTI Hybri-quick (Carl ROTH) and probed using oligonucleotide CC3271. Northern blots were revealed using the Typhoon BioMolecular Imager (Cytiva) and analyzed by ImageQuant TL (Cytiva) software.

### Cryo-EM analysis

Samples for cryo-EM analysis were prepared by mixing crude 70S ribosomes (0.54 µM), truncated *spyA-T* mRNA (2.3 µM) and PURExpress translation buffer (q.s.p 5 µL with water). QuantiFoil (300 mesh copper) R 2/1 grids were glow-discharged for 30 sec at −15 mA and 0.39 mBar using a PELCO easiGLOW device. Sample vitrification was performed at 20°C and 90% humidity with an EM GP2 (Leica) plunger. The sample (4 µL) was applied to the grids for 1 sec prior to blotting for 1.2 sec using the sensor option and was immediately plunge-frozen in liquid ethane. Grids were stored in liquid nitrogen until screening and data collection. The dataset was collected on a 200 kV Glacios (Thermo Fisher) microscope equipped with a Falcon 4 detector. Image processing was performed using cryoSPARC.

## Acknowledgments

We thank lab members for helpful discussions. We are grateful to Dr. Heather Feaga for helpful discussions and for the gift of the HAF521 strain.

---

**Supplementary Figure 1:**
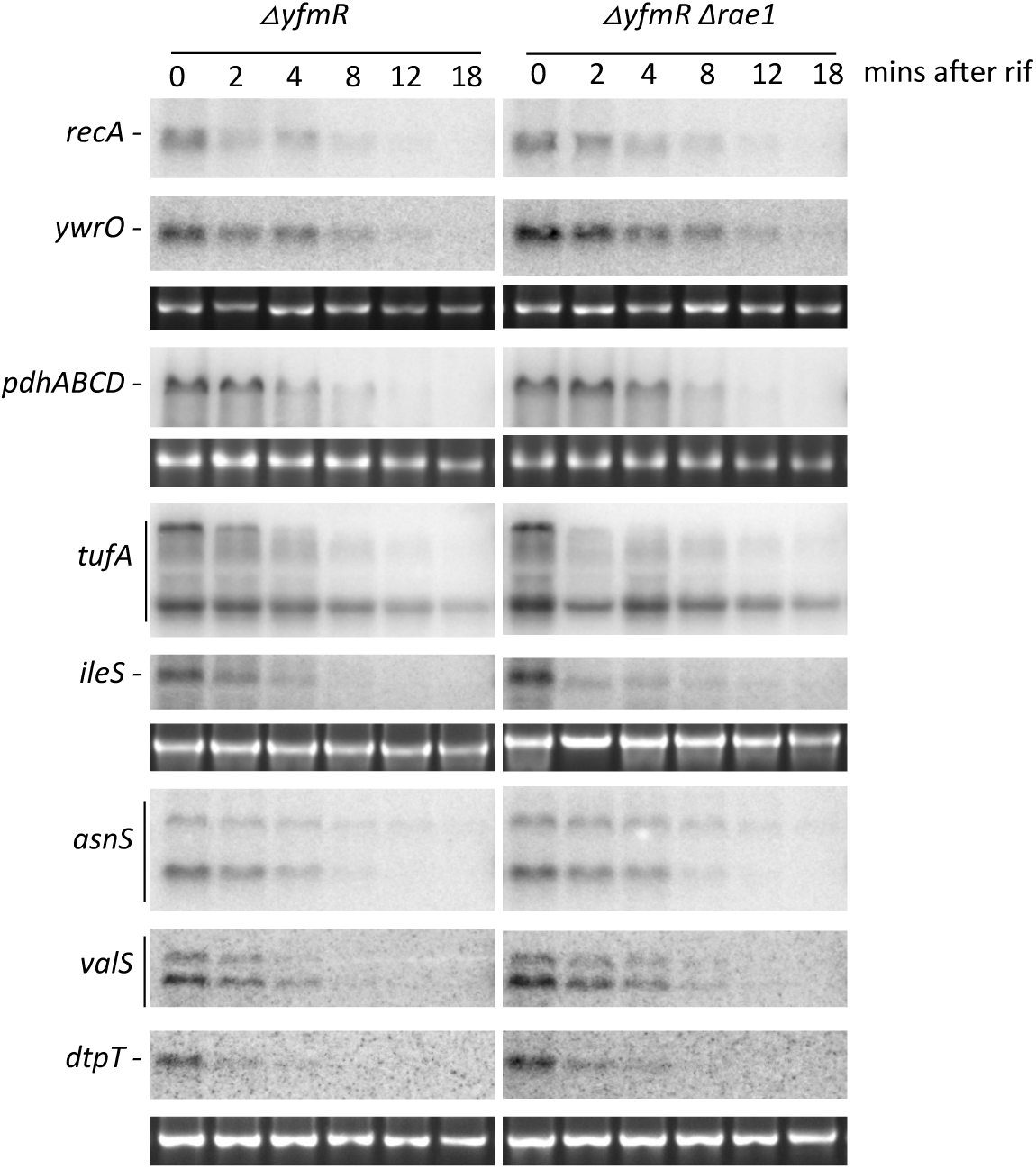
Deletion of *yfmR* has no impact on the stability of the novel Rae1 targets. The stability of mRNAs cleaved by Rae1 in absence of EF-P was analysed in △*yfmR* and △*rae1 yfmR* strains using 2 biological replicates. 16S RNA is shown as a loading control and was revealed using 1/20000 Gel Red on the agarose before transfer of the total RNA on the membrane. RNAs were probed using primers described in Supplementary Table 1. The representative northern blots of the *asnS, valS* and *dtpT* mRNAs, of *recA* and *ywrO* mRNAs, and of *tufA* and *ileS* mRNAs were probed respectively on the same northern blots.

**Supplementary Figure 2:**
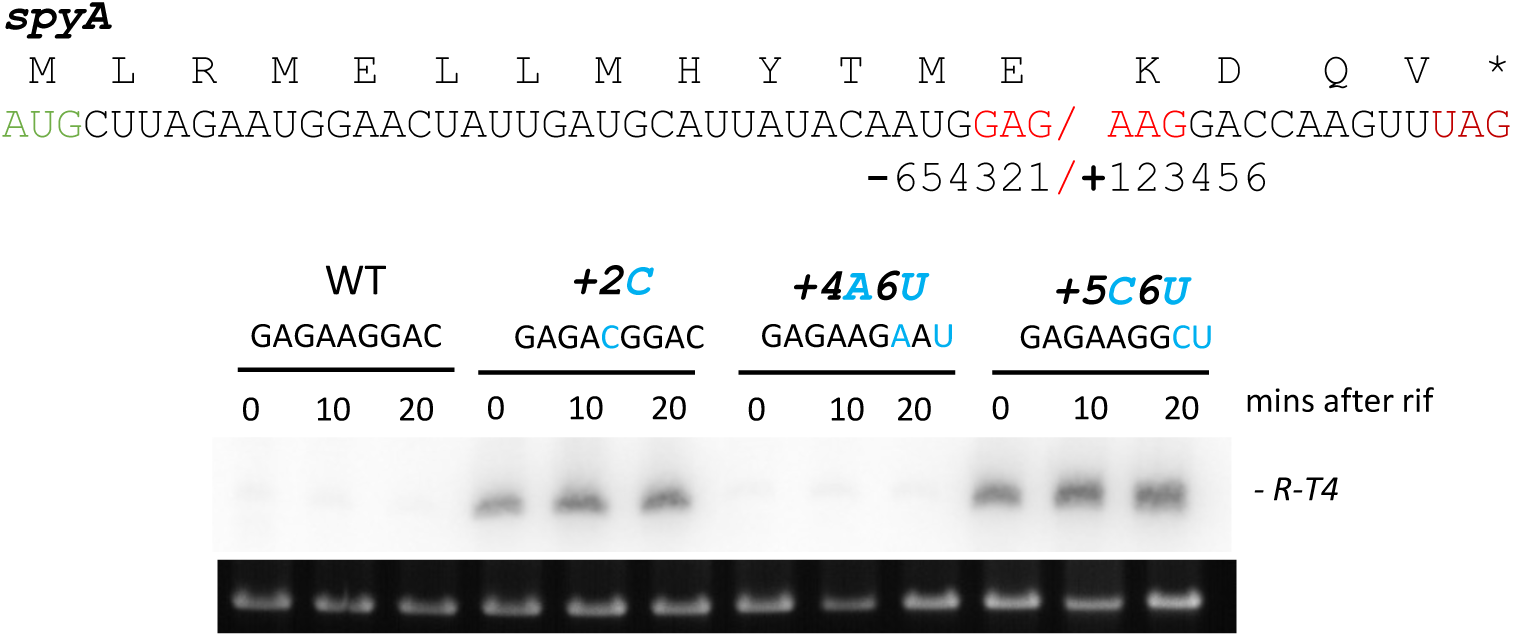
Impact of Rae1 cleavage site mutations on mRNA stability *in vivo*. *In vivo* stability of the *hbs△-spyA* constructs carrying the mutations +2C, +4A6U, and +5C6U upstream of the cleavage site (red) was quantified using 2 biological replicates. 16S RNA is shown as a loading control and was revealed using 1/20000 Gel Red staining on the agarose before transfer of the total RNA on the membrane. The R-T4 band resulting from Rae1 cleavage described in Gilet *et al.,* 2025 (Gilet et al. 2025), is indicated in black.

**Supplementary Figure 3:**
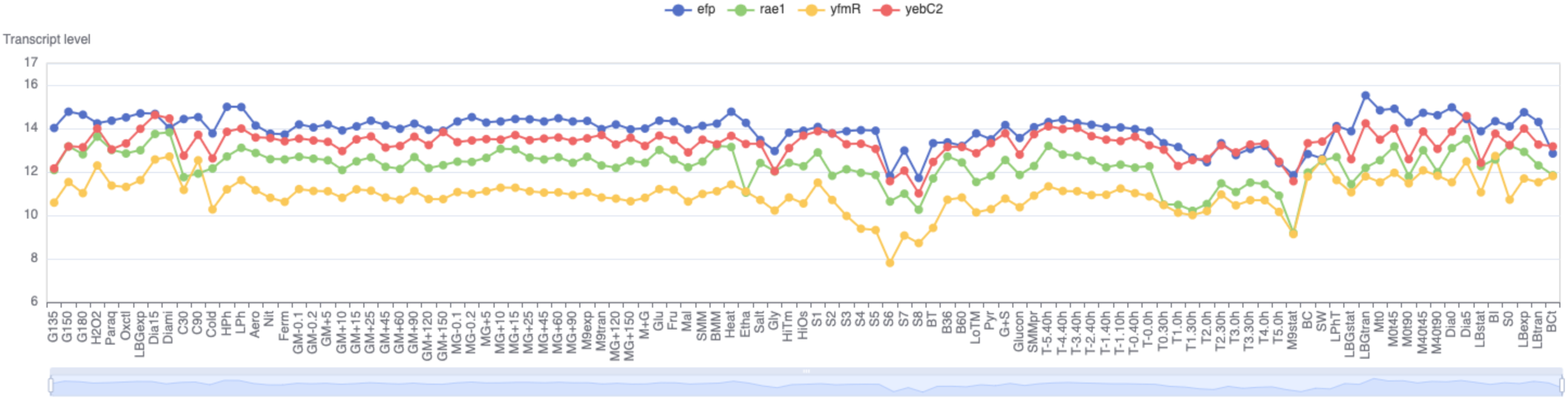
Expression profiles of *yfmR*, *yebC2*, *efp* and *rae1* mRNAs. Graphical representation from Subtiwiki (Mader et al. 2012) of the levels of the *yfmR*, *yebC2*, *efp* and *rae1* mRNAs determined by transcriptome analysis in 110 growth conditions of *B. subtilis* (Nicolas et al. 2012).

**Table S1.**
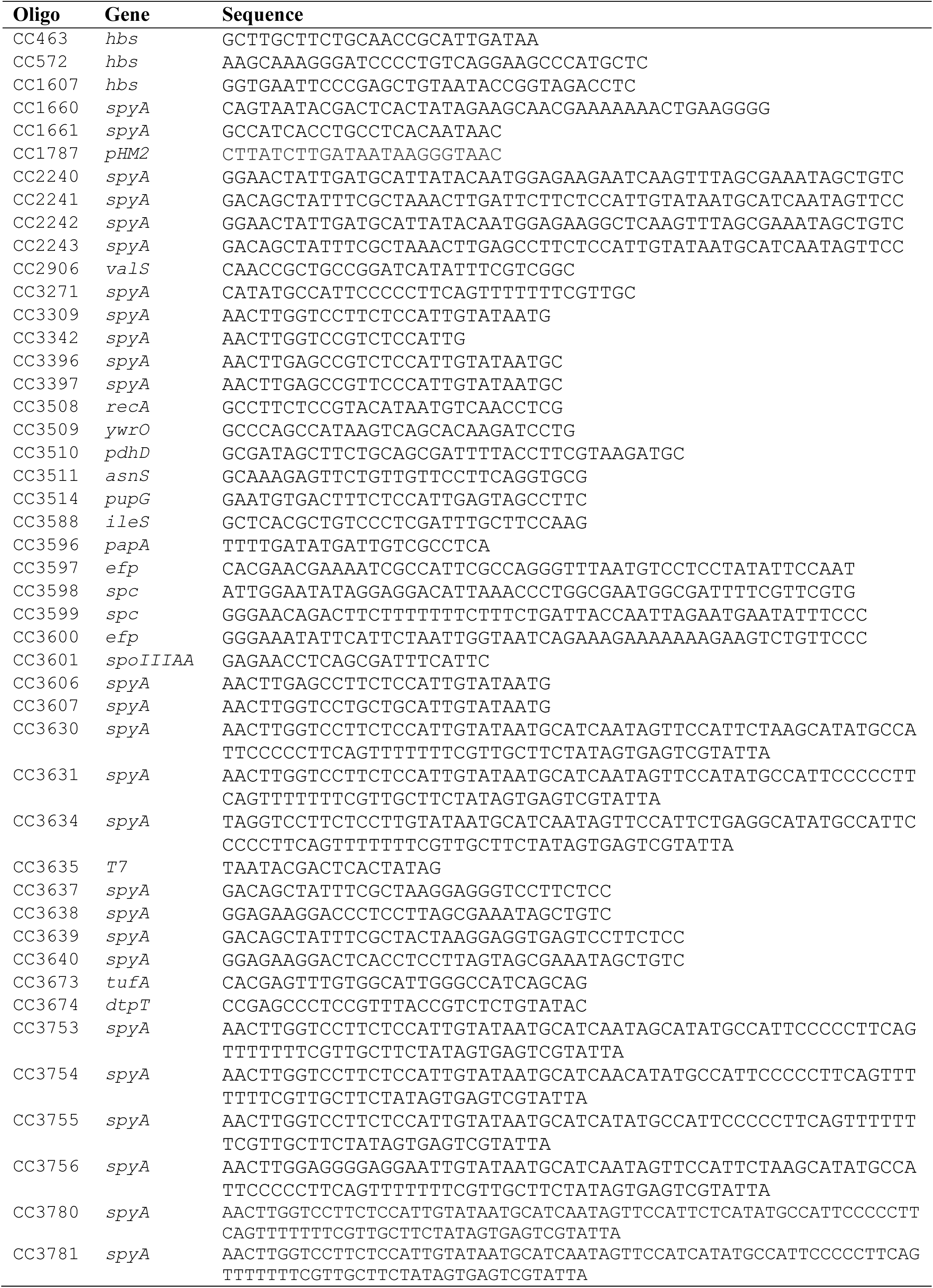
Oligonucleotides used in this study.

**Table S2.**
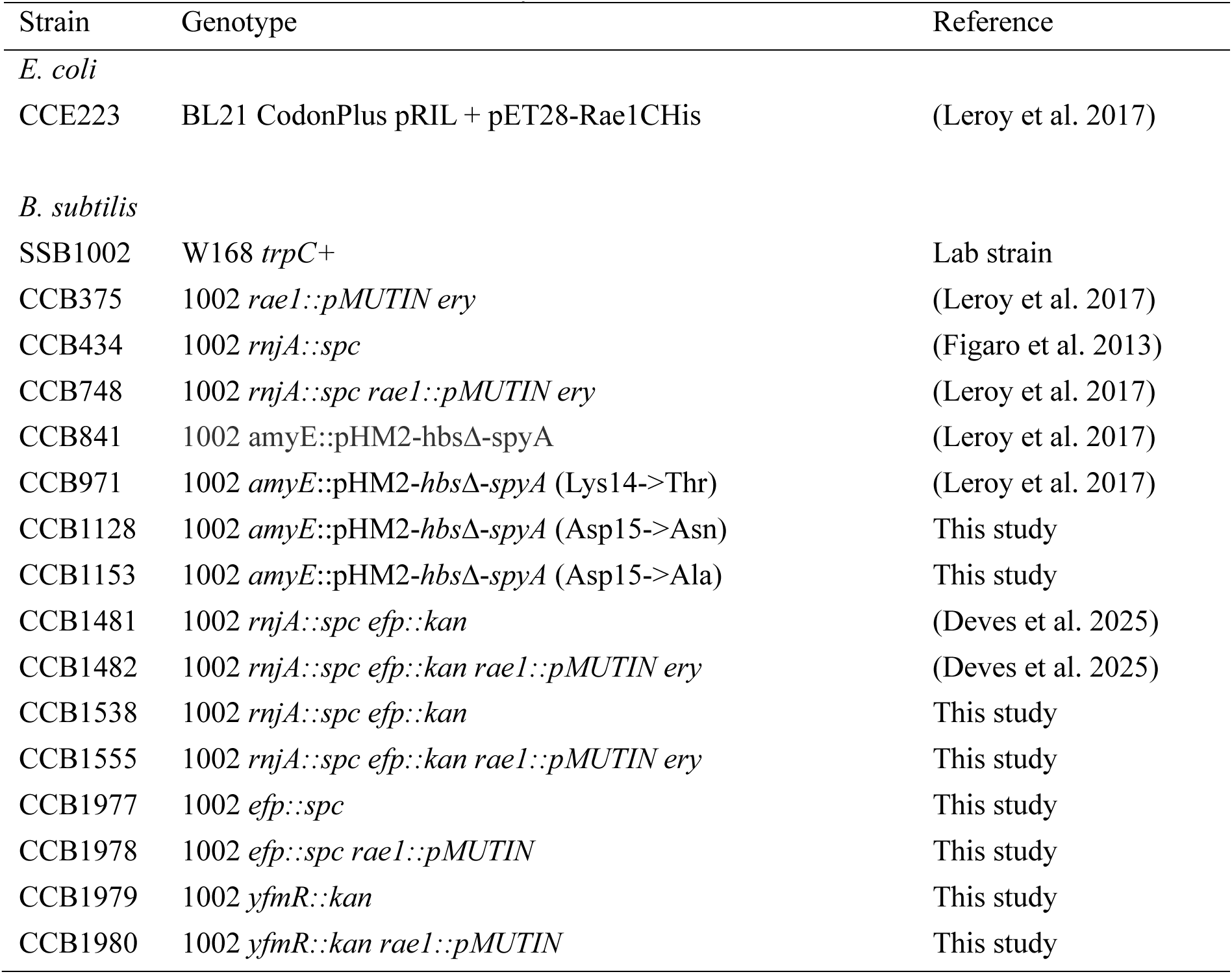
Bacterial strains used in this study.

**Table S3.**
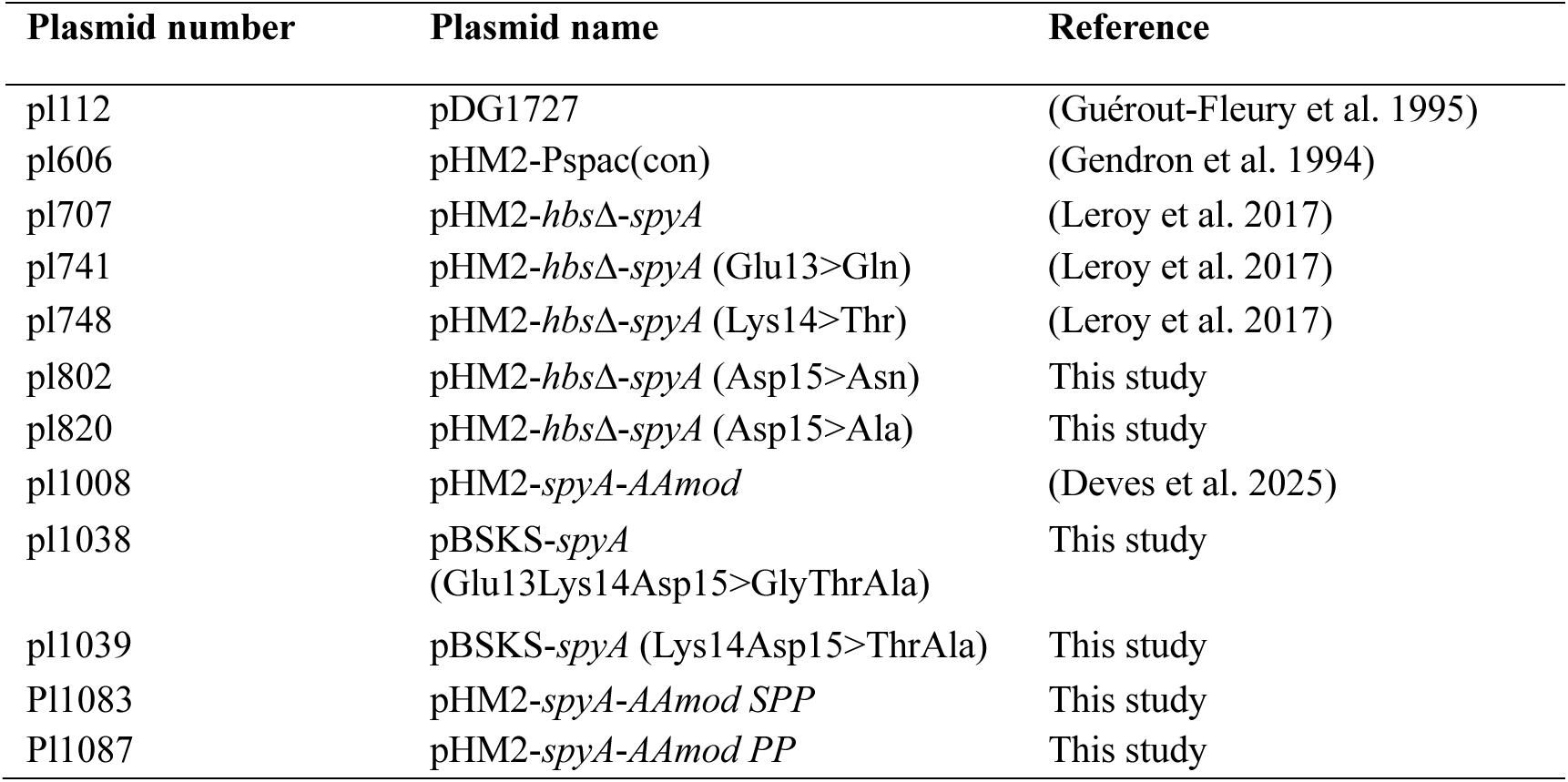
Plasmids used in this study.

| Plasmid number | Plasmid name | Reference |
| --- | --- | --- |
| pl112 | pDG1727 | (Guérout-Fleury et al. 1995) |
| pl606 | pHM2-Pspac(con) | (Gendron et al. 1994) |
| pl707 | pHM2-*hbsΔ-spyA* | (Leroy et al. 2017) |
| pl741 | pHM2-*hbsΔ-spyA* (Glu13>Gln) | (Leroy et al. 2017) |
| pl748 | pHM2-*hbsΔ-spyA* (Lys14>Thr) | (Leroy et al. 2017) |
| pl802 | pHM2-*hbsΔ-spyA* (Asp15>Asn) | This study |
| pl820 | pHM2-*hbsΔ-spyA* (Asp15>Ala) | This study |
| pl1008 | pHM2-*spyA-AAmod* | (Deves et al. 2025) |
| pl1038 | pBSKS-*spyA*<br>(Glu13Lys14Asp15>GlyThrAla) | This study |
| pl1039 | pBSKS-*spyA* (Lys14Asp15>ThrAla) | This study |
| Pl1083 | pHM2-*spyA-AAmod SPP* | This study |
| Pl1087 | pHM2-*spyA-AAmod PP* | This study |

